# Spontaneous myocarditis in mice predisposed to autoimmune disease: Including Vaccination-induced onset

**DOI:** 10.1101/2022.03.14.484354

**Authors:** Takuma Hayashi, Motoki Ichikawa, Ikuo Konishi

## Abstract

Non-obese diabetic (NOD)/ShiLtJ mice, like biobreeding rats, are used as an animal model for type 1 diabetes. Diabetes develops in NOD mice as a result of insulitis, a leukocytic infiltrate of the pancreatic islets. The onset of diabetes is associated with a moderate glycosuria and a non-fasting hyperglycemia. Previously, in non-obese diabetic (NOD)/ShiLtJ mice spontaneously developing type 1 diabetes, the possible involvement of decreased expression of LMP2/β1i, an immunoproteasome β subunit, and associated decreased expression of NF-κB1 (also called as p50) in the development of type 1 diabetes was argued between our research team and other research groups. In response to these arguments, we created NOD mice in which NF-κB1 expression is not consistently observed. Unexpectedly, most NOD *Nfκb1* homozygote mice were found to die by the 8th week of age due to the development of severe myocarditis. Furthermore, in all NOD *Nfκb1* heterozygote mice, the onset of Insulitis was observed from 4 months of age. In addition, in NOD *Nfκb1* heterozygote mice, an increase in cTnT due to vaccination with influenza or HBV vaccine was observed without gender difference. Now, we found a direct involvement of decreased expression of NF-κB1 in the development of autoimmune diseases in NOD/ShiLtJ mice. Therefore, we would like to introduce new research results on autoimmune diseases, including findings on important risk factors for the development of myocarditis observed after vaccination with mRNA-based COVID-19 vaccine.

## Introduction

Vaccination against coronavirus infection disease-2019 (COVID-19) provides clear public health benefits; however, it also carries potential risks. Recent clinical studies have revealed that the risk of myocarditis after vaccination with mRNA-based COVID-19 vaccines (BNT162b2 or mRNA-1273) increased in multiple age groups of both men and women, remarkably after the second dose in men aged 12–24 years (1). Previous clinical studies have not revealed risk factors for the development of myocarditis or endocarditis caused by mRNA-based COVID-19 vaccines. Vaccination was believed to increase the risk for autoimmune diseases, but no medical evidence has been reported. Thus, the risks and outcomes of myocarditis after mRNA-based COVID-19 vaccination are unclear.

Nuclear factor-kappa B (NF-κB) is an important transcription factor in cells that express cytokines, chemokines, growth factors, cell adhesion molecules, and some acute-phase proteins in normal and various disease states (2). NF-κB1 or NF-κB2 is bound to Rel Protooncogene (REL), RELA, or RELB to form the NF-κB complex. The transgenic mice that lack the *Nfκb1* (also as called p50) subunit of NF-κB did not show developmental abnormalities but exhibit nonspecific responses to infection, i.e., autoimmunity (3). Further studies have also suggested that NF-κB1 may be involved in the development of inflammation or autoimmune disease (4,5). Clinical studies have provided initial evidence that the −94delATTG polymorphism, which results in the defected expression of NF-κB1, may have functional attributes and appears to be an important risk factor for ulcerative colitis, an immune-mediated, complex genetic disorder (6,7). On the contrary, new mutations, e.g., in NF-κB1, PI3Kδ, PI3KR1, and PKCδ, leading to the clinical picture of common variable immune deficiency, have been shown to increasingly associate with autoimmune diseases (8,9).

We created NOD mice in which NF-κB1 expression is not consistently observed. Unexpectedly, most NOD *Nfκb1* homozygote mice were found to die by the 8th week of age due to the development of severe myocarditis. The incidence of myocarditis was 95% in males and 75% in females, with gender differences. Furthermore, in all NOD *Nfκb1* heterozygote mice, the onset of Insulitis was observed from 4 months of age. However, the incidence of myocarditis in NOD *Nfκb1* heterozygote mice was low in both males and females (15% for males and 5% for females). From the results of our research this time, it was clarified that the expression status of NF-κB1 is involved in the onset of diabetes in NOD mice. In addition, in F15-NOD *Nfκb1* heterozygote mice, an increase in serum cardiac troponin T (cTnT), which is a risk factor for myocarditis, due to vaccination with influenza or HBV vaccine was observed without gender difference. The F15-NOD *Nfκb1* heterozygote and homozygote mice, which are considered at risk of developing myocarditis, were found to have increased risk of myocarditis after vaccination with influenza or HBV vaccine. Owing to the limited data on the efficacy and safety of mRNA-based COVID-19 vaccines in patients with autoimmune diseases including collagen and rheumatic diseases, mRNA-based COVID-19 vaccination for patients with autoimmune diseases or people with allergic predisposition should be carefully considered.

## Materials and Methods

### 1. Animals

D2.129P2 (B6)-*Nfκb1* homozygote mice were purchased from JAX ^®^ Mice and Services at The Jackson Laboratory (Bar Harbor, ME, USA). NOD/ShiLtJ mice were purchased from CLEA Japan, Inc. (Meguro, Tokyo, Japan). F15 NOD *Nfκb1* heterozygote was created by back crossing D2.129P2 (B6)-*Nfκb1* homozygote male mice with NOD/ShiLtJ female mice or D2.129P2 (B6)-*Nfκb1* homozygote female mice with NOD/ShiLtJ male mice (F15 means backcrossed more than 15 times). In addition, we crossed F15 NOD *Nfκb1* heterozygote males and females to create F15 NOD *Nfκb1* homozygote.

### 2. M-mode echocardiography examination

To determine the disease severity of myocarditis, M-mode echocardiography measurements obtained from mice was performed by SonoSite M using standard procedure (FUJIFILM SonoSite, Inc., Minato-ku, Tokyo, Japan). Briefly, mice were anesthetized with ether and examined by M-mode echocardiography.

### 3. Flow cytometry

One million splenocytes obtained from D2.129P2 (B6)-*Nfκb1* homozygote mice, NOD/ShiLtJ mice, and F15 NOD *Nfκb1* heterozygote were suspended in biotin-free RPMI containing 0.1% azide and 3% FCS and surface stained in 96-well plates with the 10–3.6 PE (anti-I-Ag7) (BD PharMingen, Franklin Lakes, NJ, USA), which is class II major histocompatibility complex (MHC) haplotype for NOD/ShiLtJ mice.

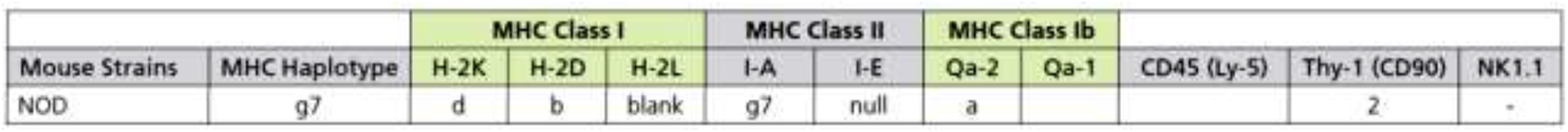

### 4. Staining and Immunohistochemistry (IHC)

IHC staining for CD3 was performed on serial heart sections and pancreatic sections obtained from F15 NOD *Nfκb1* wild type mice, F15 NOD *Nfκb1* heterozygote mice, and F15 NOD *Nfκb1* homozygote mice (Supplementary material materials and methods *1.1*). The monoclonal antibody for mouse CD3 (17A2, 1:200) was purchased from Thermo Fisher Scientific (Waltham, MA, USA). IHC was performed using the avidin–biotin complex method, as described previously [1,2]. Hematoxylin and Eosin (H.E.) staining was performed by standard procedure.

### 5. cardiac troponin T (cTnT) ELISA and myocarditis scoring

To correlate serum cardiac troponin T (cTnT) elevations with the presence and severity of myocarditis, blood was exclusively obtained at autopsy. Briefly, mice were anesthetized with ether and bled retro-orbitally. cTnT ELISA: Serum cTnT levels were determined with mouse cTnT ELISA Kit (CUSABIO TECHNOLOGY LLC, Houston, TX, USA) according to the manufacturer’s instructions.

### 6. Ethical approval and consent to participate

This study was reviewed and approved by the Central Ethics Review Board of the National Hospital Organization Headquarters in Japan (Tokyo, Japan) and Shinshu University (Nagano, Japan). The exact date when the ethical approval was obtained was August 17, 2019. The code number of the ethical approval was NHO H31-02.

Details of materials and methods are indicated in Supplementary Data sets opened online.

## Result

To understand the risks and consequences of myocarditis after COVID-19 vaccination, we examined an experimental system using small animals. Since autoimmune diseases are a polygenetic disorder, it is difficult to spontaneously induce an autoimmune disease in genetically modified mice. Therefore, we investigated changes in the symptoms of an autoimmune disease depending on the expression status of NF-κB1 in mice with a gene background of non-obese diabetic (NOD)/ShiLtJ, which are established as model mice for type 1 diabetes, an autoimmune disease (10).

The F15-NOD *Nfκb1* heterozygote was created by back-crossing D2.129P2 (B6)-*Nfκb1* homozygote mice with NOD/ShiLtJ mice 15 times (Extended Data item S.Figure 1). Additionally, we crossed F15-NOD *Nfκb1* heterozygote male and female mice to create the F15-NOD *Nfκb1* homozygote. Accordingly, approximately 30% of all infant mice born from the same F15-NOD *Nfκb1* heterozygote mother mice died by 9 weeks of age. Therefore, we performed genotyping on all infant mice. As a result, all the dead infant mice were F15-NOD *Nfκb1* homozygote. Histopathological examination and M-mode echocardiography measurements revealed myocarditis as the cause of death in both male and female infant mice by 9 weeks of age (Figure 1, Extended Data Item S.Figure 2). In NOD/ShiLtJ mice, the incidence of Insulitis in females is clearly higher than in male mice. However, Insulitis was observed in all F15 NOD *Nfκb1* heterozygote mice (Extended Data Item Table 1). The expression of *Nfκb1* was found to be involved in the onset of Insulitis. Based on the pathological diagnosis tests and results of serum cardiac troponin T (cTnT), a small part of infant mice with F15-NOD *Nfκb1* heterozygote are suspected to have mild myocarditis but matured like the F15-NOD *Nfκb1* wild-type (wt) (Figure 1, Figure 2).

**Figure 1.**
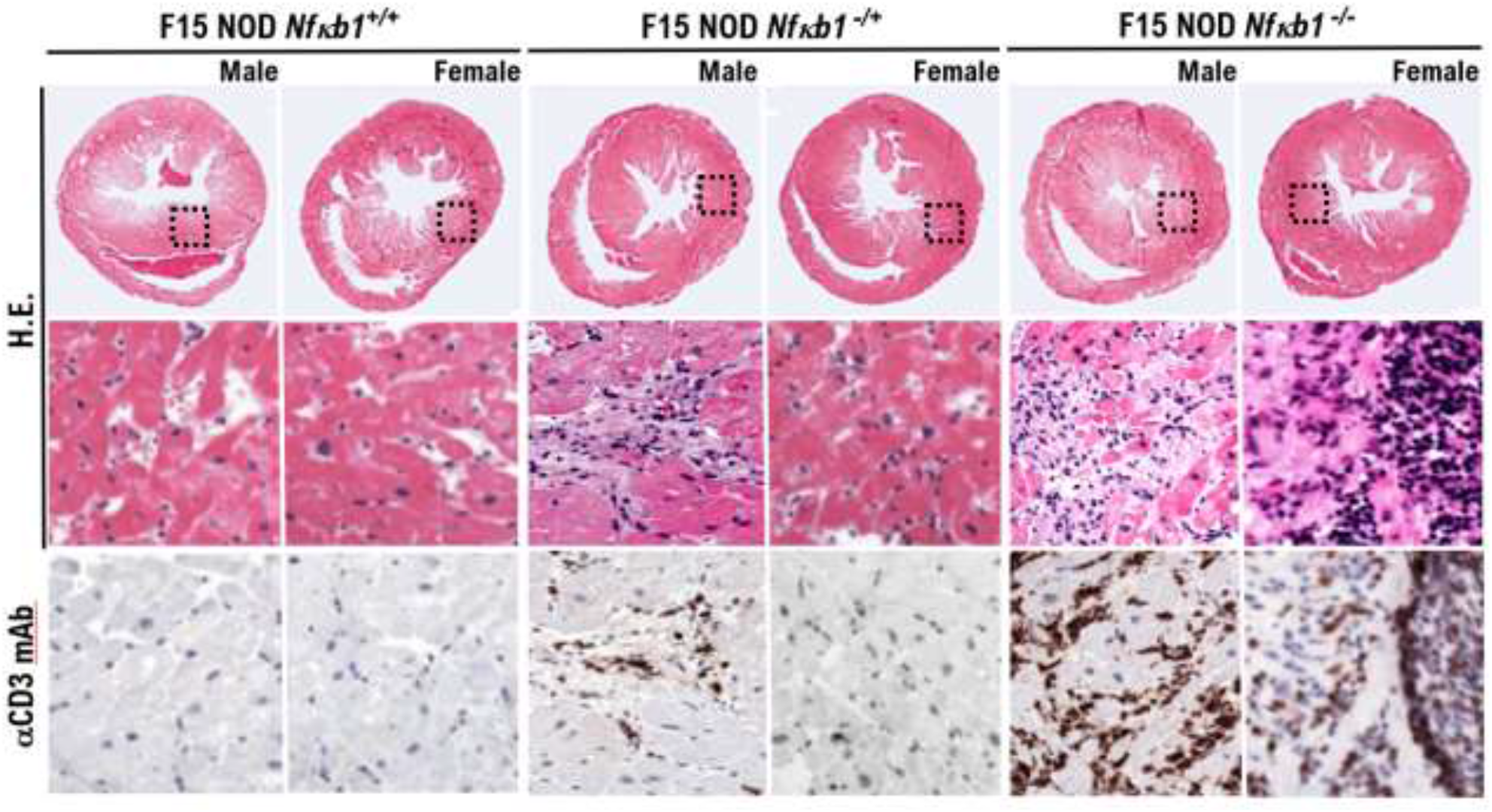
Spontaneous onset of myocarditis in F15-NOD *Nfκb1* heterozygote and homozygote male and female infant mice by 9 weeks of age. F15 hybrids of *Nfκb1* heterozygote (−/+) and homozygote (−/−) and wild-type (+/+) mice with non-obese diabetic/ShiLtJ background. Histopathological examination revealed myocarditis as the cause of death in both male and female infant mice by 9 weeks of age. Representative hematoxylin and eosin-stained tissue sections and immunohistochemically stained tissue sections with anti-CD3 monoclonal antibody of the hearts obtained from both male and female infant mice by aged 4–8 weeks. The immunohistology images show diffuse infiltration of CD3+ T cells, as shown by anti-CD3 antibody staining (brown) in the lower panels.

**Figure 2.**
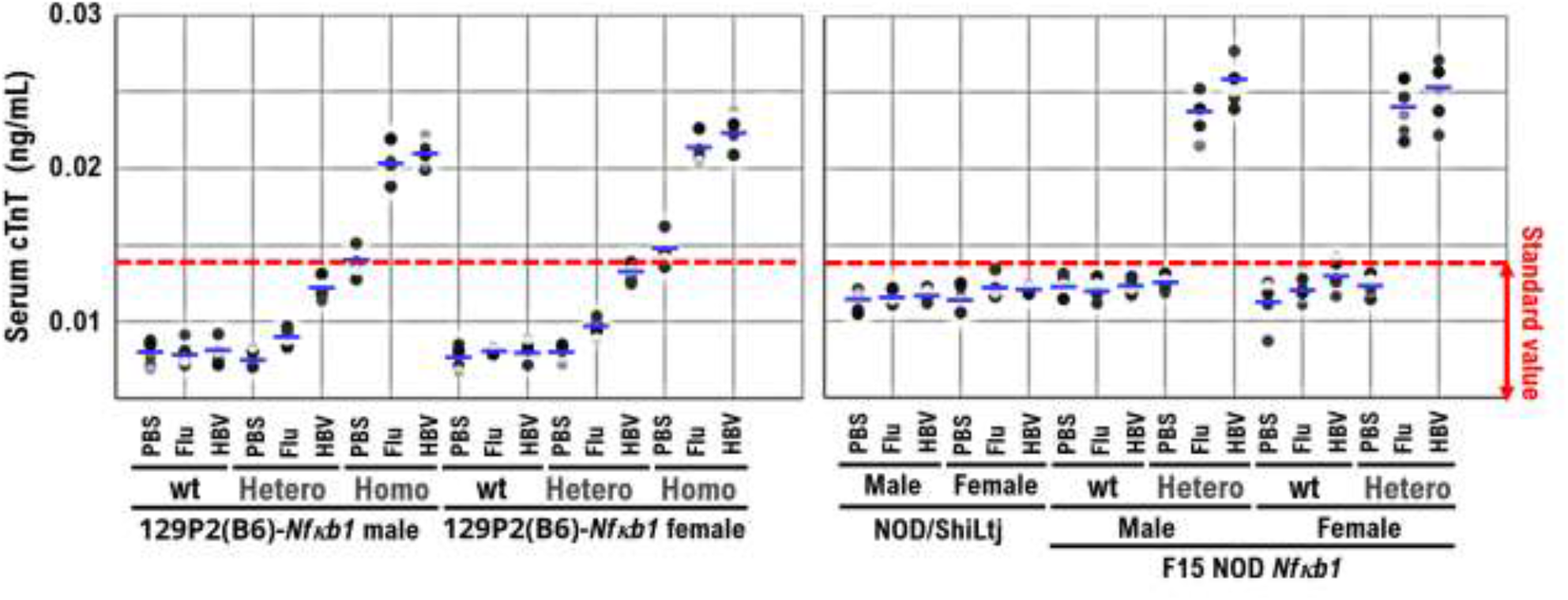
Increased serum cardiac troponin T (cTnT) concentration in 129P2 (B6)-*Nfκb1* homozygote mice and F15-NOD *Nfκb1* heterozygote mice after influenza or HBV vaccination. PBS (control), influenza vaccine, or HBV vaccine was inoculated into the left thigh muscle of 10-week-old 129P2 (B6)-*Nfκb1* wild-type (wt), 129P2(B6)-*Nfκb1* heterozygote, 129P2(B6)-*Nfκb1* homozygote, NOD/ShiLtj, F15-NOD *Nfκb1* wt, and F15-NOD *Nfκb1* heterozygote mice. Influenza or HBV vaccine was administered to mice by diluting the stock solution five-fold with PBS and inoculating 100 μL. In 129P2 (B6)-*Nfκb1* homozygote and F15-NOD *Nfκb1* heterozygote mice inoculated with influenza or HBV vaccine, the serum concentration of cTnT was ≥0.02 ng/mL. No gender difference was observed in the increase in serum cTnT concentration after each vaccination. The standard serum cTnT concentration is ≤0.014 ng/mL. In clinical practice, myocarditis is diagnosed when the serum cTnT concentration is ≥0.2 ng/mL. Each mouse group consisted of five mice.

Based on these findings, F15-NOD *Nfκb1* heterozygote and homozygote mice are at risk of developing myocarditis. Therefore, we investigated the value of cTnT, which is a risk factor for myocarditis due to the inoculation with influenza vaccine (influenza HA vaccine “KMB”) or hepatitis B virus (HBV) vaccine (Bimmugen) in NOD/ShiLtj mice and *Nfκb1* genetically modified 129P2 (B6) mice and F15-NOD mice. Consequently, in 129P2 (B6) *Nfκb1* homozygote mice, an increase in cTnT (>0.014 ng/mL) due to influenza or HBV vaccine was observed without gender difference (Figure 2). In 129P2 (B6) wt and 129P2 (B6) *Nfκb1* heterozygote mice, the cTnT value after inoculation with influenza or HBV vaccine was below the standard (0.014 ng/mL) without gender difference (Figure 2). For NOD/ShiLtj and F15-NOD *Nfκb1* wt mice, the median cTnT value for Phosphate buffered saline (PBS), influenza vaccine, or HBV vaccine ranged from 0.012 to 0.014 ng/mL. In F15-NOD *Nfκb1* heterozygote mice, an increase in cTnT (>0.02 ng/mL) due to vaccination with influenza or HBV vaccine was observed without gender difference (Figure 2). The F15-NOD *Nfκb1* heterozygote and homozygote mice, which are considered at risk of developing myocarditis, were found to have increased risk of myocarditis after vaccination with influenza or HBV vaccine.

## Discussion

Five decades ago, a man in his 20s developed myocarditis after inoculation with smallpox vaccine created by the attenuated vaccinia virus (11). Since then, myocarditis onset has been sporadically reported after various vaccinations. Since the onset of myocarditis has been observed following vaccination with attenuated virus or other main components, the etiology of myocarditis observed after inoculation with mRNA-based COVID-19 vaccines is not the genome SARS-CoV-2 mRNA. The described myocarditis cases usually occurred 10–14 days after a primary vaccination for smallpox. Furthermore, in many cases, the onset of myocarditis was observed within 5 days after inoculation with mRNA-based COVID-19 vaccines (12). This time course suggests an allergic mechanism, possibly due to the formation of immune complexes. An autoimmune mechanism was also proposed for cardiac-related adverse reactions following human papillomavirus (HPV) vaccination (13).

Furthermore, the risk for spontaneous onset of myocarditis was found to be higher in male F15-NOD *Nfkb1* homozygote mice than in female F15-NOD *Nfkb1* homozygote mice (Extended Data Item Table 1). This supports the finding that the prevalence of myocarditis after inoculation with mRNA-based COVID-19 vaccines is higher in men than in women, as evidenced in clinical practice. Further studies with these genetically modified mice may reveal the pathogenic mechanism, including gender differences in myocarditis after inoculation with mRNA-based COVID-19 vaccines. Owing to the limited data on the efficacy and safety of mRNA-based COVID-19 vaccines in patients with autoimmune diseases including collagen and rheumatic diseases, mRNA-based COVID-19 vaccination for patients with autoimmune diseases or people with allergic predisposition should be carefully considered.

## Supplementary Materials

These are available online.

## Author Contributions

TH, and MI performed most of the clinical work and coordinated the project. TH and MI conducted the diagnostic pathological studies. TH conceptualized the study and wrote the manuscript. TH, MI, and IK and carefully reviewed this manuscript and commented on the aspects of medical science. IK shared information on clinical medicine and oversaw the entirety of the study.

## Acknowledgments

We thank all medical staff for providing medical care to this patient at the National Hospital Organization Kyoto Medical Center. We appreciate Crimson Interactive Japan Co., Ltd., for revising and polishing our manuscript. This clinical research was performed with research funding from the following: Japan Society for Promoting Science for TH (Grant No. 19K09840), for KA (No. 20K16431), and START-program Japan Science and Technology Agency for TH (Grant No. STSC20001), and the National Hospital Organization Multicenter clinical study for TH (Grant No. 2019-Cancer in general-02).

## References

1. Oster ME, et al. JAMA. 327, 331–340 (2022).

2. Barnes, P. J., Karin, M. New Eng. J. Med. 336, 1066–1071 (1997).

3. Sha, W. C., Liou, H.-C., Tuomanen, E. I., Baltimore, D. Cell 80, 321–330 (1995).

4. Mariani S. et al. Nature Medicine 6(1), 27 (2000).

5. Hayashi T. et al. Nature Medicine 6(10), 1065–1066 (2000).

6. Hegazy, D.M et al. Genes Immun. 2, 304–308 (2001).

7. Karban AS et al Human Molecular Genetics, 13(1), 35–45 (2004)

8. Fliegauf, M., et al. Am. J. Hum. Genet. 97, 389–403 (2015).

9. Leavis H. et al. Curr Rheumatol Rep. 21(10), 55 (2019).

10. Delovitch TL, Singh B. Immunity 7, 727–738 (1997).

11. Matthews A.W., Griffiths I.D. Br Heart J. 36(10), 1043–1045 (1974).

12. Montgomery J, et al. JAMA Cardiol. 6(10), 1202–1206 (2021).

13. Ryabkova VA, et al. Autoimmun Rev. 18(4), 415–425 (2019).

