## Supplemental 1 for "Spontaneous myocarditis in mice predisposed to autoimmune disease: Including Vaccination-induced onset"

### Supplementary material

### 1. Materials and Methods

#### *1.1. Animals*

D2.129P2 (B6)-*Nfκb1* homozygote mice were purchased from JAX ® Mice and Services at The Jackson Laboratory (Bar Harbor, ME, USA). NOD/ShiLtJ mice were purchased from CLEA Japan, Inc. (Meguro, Tokyo, Japan). F15 NOD *Nfκb1* heterozygote was created by back crossing D2.129P2 (B6)-*Nfκb1* homozygote male mice with NOD/ShiLtJ female mice or D2.129P2 (B6)-*Nfκb1* homozygote female mice with NOD/ShiLtJ male mice (F15 means backcrossed more than 15 times). In addition, we crossed F15 NOD *Nfκb1* heterozygote males and females to create F15 NOD *Nfκb1* homozygote. All experiments were performed using age and sex-matched groups. Mice were maintained in specific pathogen-free facilities at Shinshu University (Matsumoto, Nagano, Japan). All of the study protocols were approved by the Animal Care and Use Committee of the Shinshu University and National Hospital Organization Kyoto Medical Center (Approved number 2016-3, 2019-03). Experiments were performed according to the official rules formulated in the Japanese law on the care and use of experimental animals.

The genotyping protocol(s) presented here have been optimized for reagents and conditions used by The Jackson Laboratory (JAX). The genotyping analyses were performed using primer sets as followings, Common 5’-GCA AAC CTG GGA ATA CTT CAT GTGACT AAG-3’, wild type 5’-ATA GGC AAG GTC AGA ATG CAC CAGAAG TCC-3’, mutant 5’-AAA TGT GTC AGT TTC ATA GCC TGAAGA ACG-3’. The detail of genotyping protocol by PCR is described in following JAX website, https://www.jax.org/Protocol?stockNumber=002849&protocolID=19175

#### *1.2. M-mode echocardiography examination.*

To determine the disease severity of myocarditis, M-mode echocardiography measurements obtained from mice was performed by SonoSite M using standard procedure (FUJIFILM SonoSite, Inc., Minato-ku, Tokyo, Japan). Briefly, mice were anesthetized with ether and examined by M-mode echocardiography.

#### *1.3. Flow cytometry.*

One million splenocytes obtained from D2.129P2 (B6)-*Nfκb1* homozygote mice, NOD/ShiLtJ mice, and F15 NOD *Nfκb1* heterozygote were suspended in biotin-free RPMI containing 0.1% azide and 3% FCS and surface stained in 96-well plates with the 10–3.6 PE (anti-I-Ag7) (BD PharMingen, Franklin Lakes, NJ, USA), which is class II major histocompatibility complex (MHC) haplotype for NOD/ShiLtJ mice. The splenocytes were washed no fewer than two times before the addition of the secondary reagent. All samples were analyzed on a FACSCalibur flow cytometer (Becton Dickinson, Mountain View, CA, USA) using CellQuest software (Becton Dickinson).

| | | MHC Class I | | | MHC Class II | | MHC Class Ib | | | | |
|---|---|---|---|---|---|---|---|---|---|---|---|
| Mouse Strains | MHC Haplotype | H-2K | H-2D | H-2L | I-A | I-E | Qa-2 | Qa-1 | CD45 (Ly-5) | Thy-1 (CD90) | NK1.1 |
| NOD | g7 | d | b | blank | g7 | null | a | | | 2 | - |

https://tools.thermofisher.com/content/sfs/brochures/Mouse_Haplotype_Table.pdf

***1.4. Staining and Immunohistochemistry (IHC).***

IHC staining for CD3 was performed on serial heart sections and pancreatic sections obtained from F15 NOD *Nfκb1* wild type mice, F15 NOD *Nfκb1* heterozygote mice, and F15 NOD *Nfκb1* homozygote mice (Supplementary material materials and methods *1.1*). The monoclonal antibody for mouse CD3 (17A2, 1:200) was purchased from Thermo Fisher Scientific (Waltham, MA, USA). IHC was performed using the avidin–biotin complex method, as described previously [1,2]. Hematoxylin and Eosin（H.E.）staining was performed by standard procedure. Briefly, one representative 5-mm tissue section was cut from a paraffin-embedded sample of heart and pancreatic tissues obtained from F15 NOD *Nfκb1* wild type mice, F15 NOD *Nfκb1* heterozygote mice, and F15-NOD *Nfκb1* homozygote mice.

Next, the sections were incubated with a biotinylated secondary antibody (Dako, DK-2600 Glostrup, Denmark) and then incubated with a streptavidin complex (Dako). The completed reaction was developed by 3, 39-diaminobenzidine, and the slide was counterstained with hematoxylin. Normal myometrium portions in the specimens were positive controls. The negative controls comprised tissue sections incubated with normal rabbit IgG instead of the primary antibody. The expression of CD3 is indicated by brown 3,3'-Diaminobenzidine, tetrahydrochloride (DAB) staining. Normal rabbit antiserum was a negative control for the primary antibody. The entire brown DAB-stained tissue was scanned with a digital microscope BZ-X800 (Keyence, Osaka, Osaka, Japan). Hearts and pancreas were then removed from mice, fixed in 4% formalin, and paraffin embedded. Sections were obtained at eight different levels and stained with H.E. by standard procedure. Brown dots indicate the lymphocytes expressing murine CD3. Shinshu University approved these experiments according to internal guidelines (approval no. M192).

***1.5. cardiac troponin T (cTnT) ELISA and myocarditis scoring.***

To correlate serum cardiac troponin T (cTnT) elevations with the presence and severity of myocarditis, blood was exclusively obtained at autopsy. Briefly, mice were anesthetized with ether and bled retro-orbitally. Alternatively, blood was obtained by cardiac puncture. Serum samples were stored individually at −70°C until use. Immediately after bleeding, mice were killed by cervical dislocation. Hearts were then removed, fixed in 4% formalin, and paraffin embedded. Sections were obtained at eight different levels and stained with H.E.. The diagnosis of myocarditis was established by the presence of an inflammatory cell infiltrate and myocyte damage. The disease severity was determined according to a previously described scoring system ranging from 0 to 4 (1 corresponds to infiltration of ≤5% of at least one histological cross section; 2, 5% to 10%; 3, 10% to 20%; and 4, >20%) (3). Differences in disease severity were analyzed by ANOVA for multiple-sample comparisons (Bonferroni).

cTnT ELISA: Serum cTnT levels were determined with mouse cTnT ELISA Kit (CUSABIO TECHNOLOGY LLC, Houston, TX, USA) according to the manufacturer’s instructions. This sandwich ELISA system is based on two mAbs, namely, a biotinylated cTnT–specific capture antibody (CSB-PA024016HA01MO, CUSABIO TECHNOLOGY LLC) that binds to streptavidin-coated plastic tubes and a horseradish peroxidase–labeled detection antibody.

***1.6. Ethical approval and consent to participate.***

This study was reviewed and approved by the Central Ethics Review Board of the National Hospital Organization Headquarters in Japan (Tokyo, Japan) and Shinshu University (Nagano, Japan). The exact date when the ethical approval was obtained was August 17, 2019. The code number of the ethical approval was NHO H31-02. The authors attended educational lectures on medical ethics in 2020 and 2021, which were supervised by the Japanese government. The completion numbers for the authors are AP0000151756, AP0000151757, AP0000151769, and AP000351128. Consent to participate was required as this research was a clinical research. Subjects signed the informed consent when they were briefed on the clinical study and agreed with contents of clinical research. The authors attended a seminar on the ethics of experimental research using small animals on July 02, 2020 and

July 20, 2021. They became familiar with the importance and ethics of animal experiments (National Hospital Organization Kyoto Medical Center and Shinshu University School of Medicine). The code number of the ethical approval for experiments with small animal was KMC R02-0702.

### References

1. Hayashi, T.; Kobayashi, Y.; Kohsaka, S.; Sano, K. The mutation in the ATP binding region of JAK1, identified in human uterine leiomyosarcomas, results in defective interferon-gamma inducibility of TAP1 and LMP2. *Oncogene* 2006, *25*, 4016-4026. PMID: 16474838 DOI: 10.1038/sj.onc.1209434
2. Hayashi, T.; Ichimura, T.; Yaegashi, N.; Shiozawa, T.; Konishi, I. Expression of CAVEOLIN 1 in uterine mesenchymal tumors: No relationship between malignancy and CAVEOLIN 1 expression. *Biochem Biophys Res Commun.* 2015, *463(4)*, 982-987. PMID: 26072376 DOI: 10.1016/j.bbrc.2015.06.046
3. Laemmli UK. Cleavage of structural proteins during the assembles of bacterophage T4. Nature. 1970; *227*:680-685. DOI: 10.1063/1.2142136

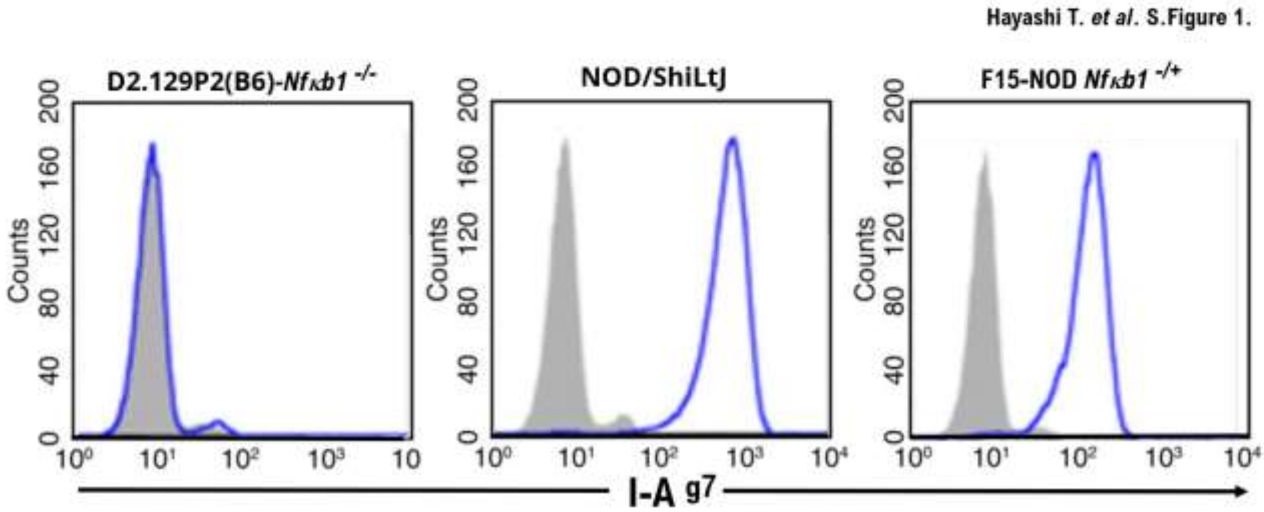


**S.Figure 1**. The expression of class II MHC haplotype I-Ag7 on splenocytes obtained from F15 NOD *Nfκb1* heterozygote mice. Fluorescence-activated cell sorter (FACS) analysis was used to assess the levels of antigen-presenting machinery by class II MHC on splenic B-cells as determined by gating on B220-positive cells in splenocytes derived from splenocytes of D2.129P2(B6)-*Nfkb1* homozygote (-/-) mice, NOD/ShiLtJ mice, and F15 NOD *Nfκb1* heterozygote (-/+) mice. B220 positive B cells were stained with anti–I-$A^{g7}$ (blue lines), respectively, and then analyzed by FACS. The gray lines indicate the isotype control for each cell line. Histograms are representative of more than three experiments in which at least three mice were analyzed. Comparison of mean fluorescence intensities among analyzed samples demonstrated $P < 0.05$ for data represented.

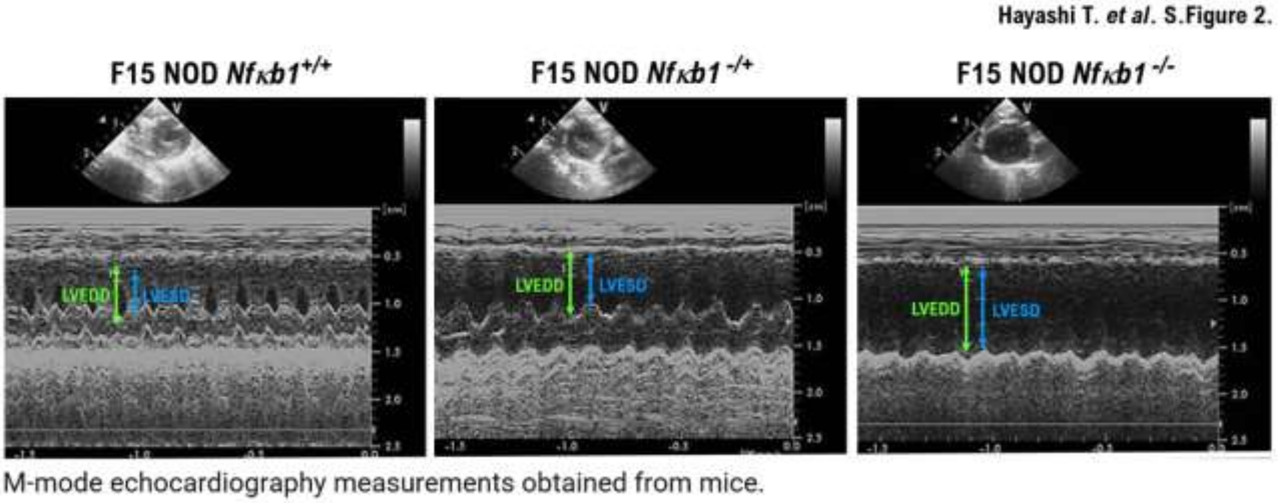


**S.Figure 2.** M-mode echocardiography measurements obtained from F15 NOD *Nfκb1* wild type (+/+) mice at 5 weeks of age, age matched F15 NOD *Nfκb1* heterozygote (-/+) mice, and F15 NOD *Nfκb1* homozygote (-/-) mice. LVEDD, left ventricle end-diastolic dimension; LVESD, left ventricle end-systolic dimension.
